# High speed multi-plane super-resolution structured illumination microscopy of living cells using an image-splitting prism

**DOI:** 10.1101/773440

**Authors:** Adrien Descloux, Marcel Müller, Vytautas Navikas, Andreas Markwirth, Robin Van den Eynde, Tomas Lukes, Wolfgang Hübner, Theo Lasser, Aleksandra Radenovic, Peter Dedecker, Thomas Huser

**Author notes:** These authors contributed equally to this work (Adrien Descloux, Marcel Müller). Corresponding authors: Aleksandra Radenovic, Peter Dedecker and Thomas Huser.

## Abstract

Super-resolution structured illumination microscopy (SR-SIM) can be conducted at video-rate acquisition speeds when combined with high-speed spatial light modulators and sCMOS cameras, rendering it particularly suitable for live cell imaging. If, however, three-dimensional (3D) information is desired, the sequential acquisition of vertical image stacks employed by current setups significantly slows down the acquisition process. In this work we present a multi-plane approach to SR-SIM that overcomes this slowdown via the simultaneous acquisition of multiple object planes, employing a recently introduced multi-plane image splitting prism combined with high-speed SR-SIM illumination. This strategy requires only the introduction of a single optical element and the addition of a second camera to acquire a laterally super-resolved three-dimensional image stack. We demonstrate the performance of multi-plane SR-SIM by applying this instrument to the dynamics of live mitochondrial network.

## 1 Introduction

Conventional fluorescence microscopy is inherently limited in its spatial resolution due to diffraction. Optical super-resolution imaging techniques allow us to overcome this limitation. For super-resolution structured illumination microscopy (SR-SIM) this is achieved by using high spatial illumination frequencies that down-modulate spatial frequencies beyond the cut-off into the passband of the microscope[1, 2, 3, 4].

Linear implementations of SR-SIM create a sinusoidal interference pattern in the sample plane, leading to a spatial resolution improvement of at best two-fold over wide-field microscopy, and additional contrast enhancement for high spatial frequencies and strong suppression of out-of-focus light by filling the missing cone of the instrument’s optical transfer function[3, 5, 6]. In return, only a small number of raw images with defined illumination patterns (9 in the standard 2D implementation, less when using advanced algorithms[7]) are needed for the image reconstruction process. The excitation powers are comparable to conventional wide-field imaging, so photo-damage can be minimized, and no special dyes (e.g. dyes that are photo-switchable or inherently blinking) are required for the linear 2x resolution enhancement. The combination of these features makes SR-SIM a very fast super-resolution imaging technique, with current instruments[8, 9] providing 2D imaging with approximately 100 nm spatial resolution in the lateral direction at video-rate speed and even faster.

Live-cell imaging is the primary domain for SR-SIM. Here, its high temporal resolution excels, especially when observing highly dynamic processes on sub-second time-scales. If, however, 3D volumetric imaging is required, current state-of-the-art SR-SIM systems slow down the acquisition rate significantly, due to the need to physically move the object through the focal plane, e.g. by piezo-translation stages. The slowdown arises not just from the repeated image acquisitions, but also from the intermediate sample movement that needs to be accounted for. Also, advanced control electronics are required if stage movement, SR-SIM illumination and camera exposure are to be synchronized precisely.

Such delays can be avoided if a full 3D volume of the sample could be acquired at once. Two different approaches to multi-plane image detection currently exist: A diffractive element, based on a phase-shifting spatial light modulator (SLM) or a lithographically-produced optical element[10, 11], can be introduced into a conjugated image plane and create almost arbitrary multi-plane detection schemes. Chromatic aberrations can be corrected using additional elements[12]. Recently, this approach was combined with a commercial SR-SIM microscope[13] and provided the first multi-plane SR-SIM images in a prototype system. Although a great demonstration of the future of SR-SIM imaging, these tests ended up being limited by the slow speed of the commercial SR-SIM platform, which was not originally conceived for video-rate super-resolution imaging. Here, we present a multi-plane imaging approach to SR-SIM that circumvents these issues and achieves high-speed imaging rates previously only available to 2D SR-SIM.

## 2 Multi-plane SR-SIM

Our work is based upon an imaging strategy for multi-plane detection that divides the detected fluorescence light into multiple image planes by introducing discrete optical pathlength differences to the image path [14, 15, 16]. This path length difference guarantees perfect object-image conjugation for 8 object planes equally displaced along the axial direction. As seen in Fig. 1, the classical 8f set-up of the detection path allows us to easily realize a telecentric instrument ensuring identical scaling of all 8 images obtained from different axially displaced object planes.

**Figure 1:**
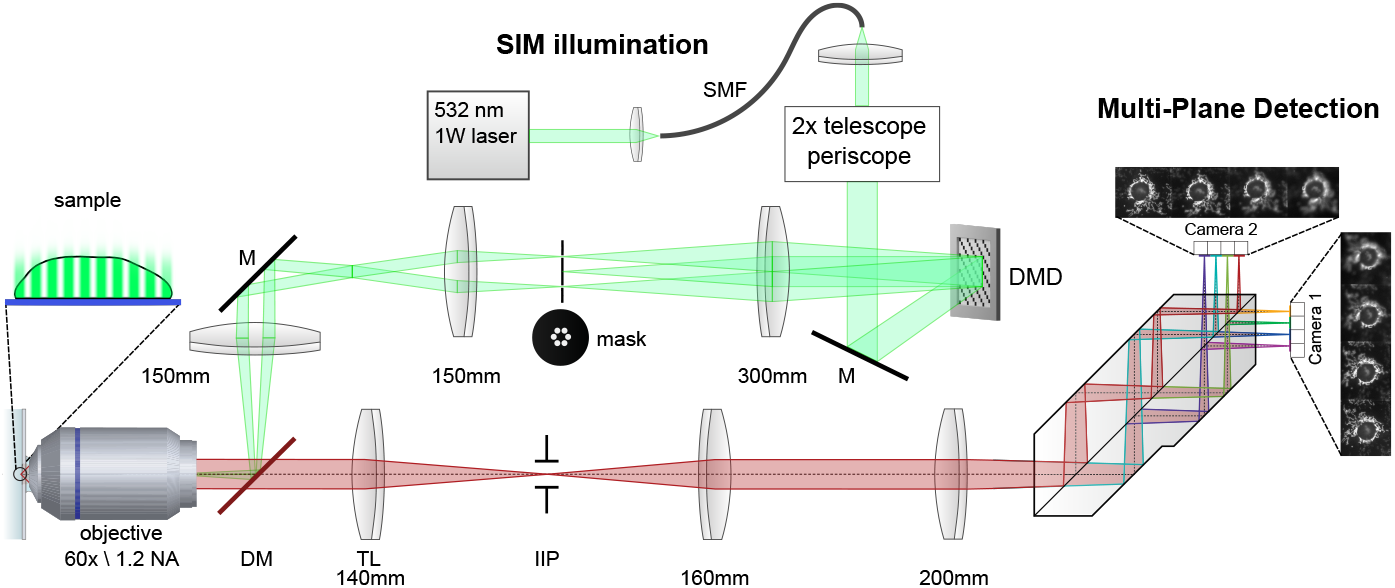
Opto-mechanical schematics of the high-speed multi-plane SR-SIM imaging system. The SR-SIM pattern is created by a 532 nm 1 W laser (Roithner) coupled into a single mode fiber for mode cleaning. The light at the distal end of the fiber is then collimated, expanded by a factor of 2 and directed onto a digital micro-mirror device (DMD), which is used as a fast, electronically controlled optical grating. A segmented aperture mask removes spurious diffraction orders, arising due to the binary nature of the DMD. A relay lens system focuses the light into the back focal plane of the objective lens (Olympus UPLSAPO 60 x/1.2 NA water immersion). Excitation and emission light is wavelength-separated by a dichroic mirror (DM). In the detection path, the tube lens (TL) forms an intermediate image (*Image Plane*), where a field stop (IIP) sets the field-of-view size and prevents overlapping signals in the multi-plane detection system. A magnification of 58.5 x is achieved, which corresponds to a projected pixel size of 111 nm for the sCMOS cameras (ORCA Flash 4.0, Hamamatsu). Lenses form a relay telescope into the main component, the precision-made multi-plane prism [16]. It provides 2 × 4 copies of the image onto 2 scientific sCMOS cameras, where each copy has a defined path-length difference and thus defined, axially offset focal plane.

While image splitting via subsequent changes in detection path length is a robust concept, its typical implementation with discrete optical elements is sensitive to thermal drifts and vibration and rather difficult to align. A recent development[16] solved this problem by integrating this approach into a single, stable, precision-made optical element, an image-splitting prism. This device allows for the simultaneous detection of up to 8 diffraction limited images obtained at equally spaced vertical planes with only minimal alignment of the overall optical set-up.

We combined our prism-based detection path with a SR-SIM excitation path based on a high-speed digital mirror device (DMD - DLP7000, Vialux) and a high-power laser (532 nm, 1W, Roithner). By utilizing two synchronized sCMOS cameras (Hamamatsu Orcaflash V4.0), we achieved high-speed volumetric imaging with 50 ms exposure time. Taking camera readout and device synchronization into account, this results in about 1.3 reconstructed SIM data sets per seconds, to image a full volume of about 40 × 40 × 2.45 *μm*^3^. Both the DMD-based spatial light modulator and the cameras could be tuned to provide even higher imaging speeds[8], if required by the application and if bright enough fluorophores are used to compensate for the lower signal to noise ratio.

In its current configuration, the SIM is operated in *2-beam* mode. Here, the ±1^st^ diffraction orders generated by the SIM grating are interfered in the sample plane to generate a lateral modulation of the excitation pattern. The 0^th^ order is blocked, so no axial modulation of the SIM pattern is introduced. This approach sacrifices the axial resolution improvement possible in full 3-beam SIM, but comes with a set of advantages. First of all, including SIM modulation along the axial direction would entail aligning each z-plane with the axial periodicity of the SIM pattern, which would take away some of the systems robustness and simplicity. Also, 15 instead of 9 raw images are required for a direct SIM reconstruction of a 3-beam signal, which impacts overall imaging speed, as well as sample photobleaching and phototoxicity. Most importantly, the optical sectioning intrinsically provided by 3-beam SIM can also be obtained in a 2-beam approach by trading in some of the lateral resolution enhancement for optical sectioning capability[5, 6]. This is achieved by using a slightly coarser SIM pattern (frequency of 1.8 *μm*^−1^) and has the additional effect of simplifying the polarization control typically necessary for SIM illumination[17]. We thus tuned the SIM system to a compromise of both optical sectioning and lateral resolution improvement, while operating at the minimum of 9 raw SIM frames to achieve high imaging speeds.

## 3 Results

We applied our multi-plane SR-SIM system to the fast imaging of mitochondrial dynamics [18, 19]. Here, SR-SIM is an enabling imaging method, as its increase in spatial resolution and inherent background suppression allows us to visualize the mitochondrial ultra-structure, the cristae, which cannot be observed with conventional wide-field resolution [20]. Furthermore, the rapid movement of mitochondria requires high imaging speed, while their 3-dimensional nature greatly benefits from acquiring signals from multiple z-planes. However, the structures are still rather “thin” compared to their length, and a coverage of 2.45 *μm* (8 planes with an axial sampling of 350 nm) as provided by our system is typically sufficient to capture the entire three-dimensional mitochondrial structure in a single shot multi-plane exposure. In this case, no axial movement of the sample is needed at all, which greatly simplifies data acquisition and decreases the acquisition time. The data processing and results are presented in Fig. 2, where our imaging was performed with 50 ms exposure time per raw data frame. As mentioned earlier, 3 angles and 3 phases are required for the 2D SIM image reconstruction process, thus reaching 770 ms acquisition time for the full imaged volume.

**Figure 2:**
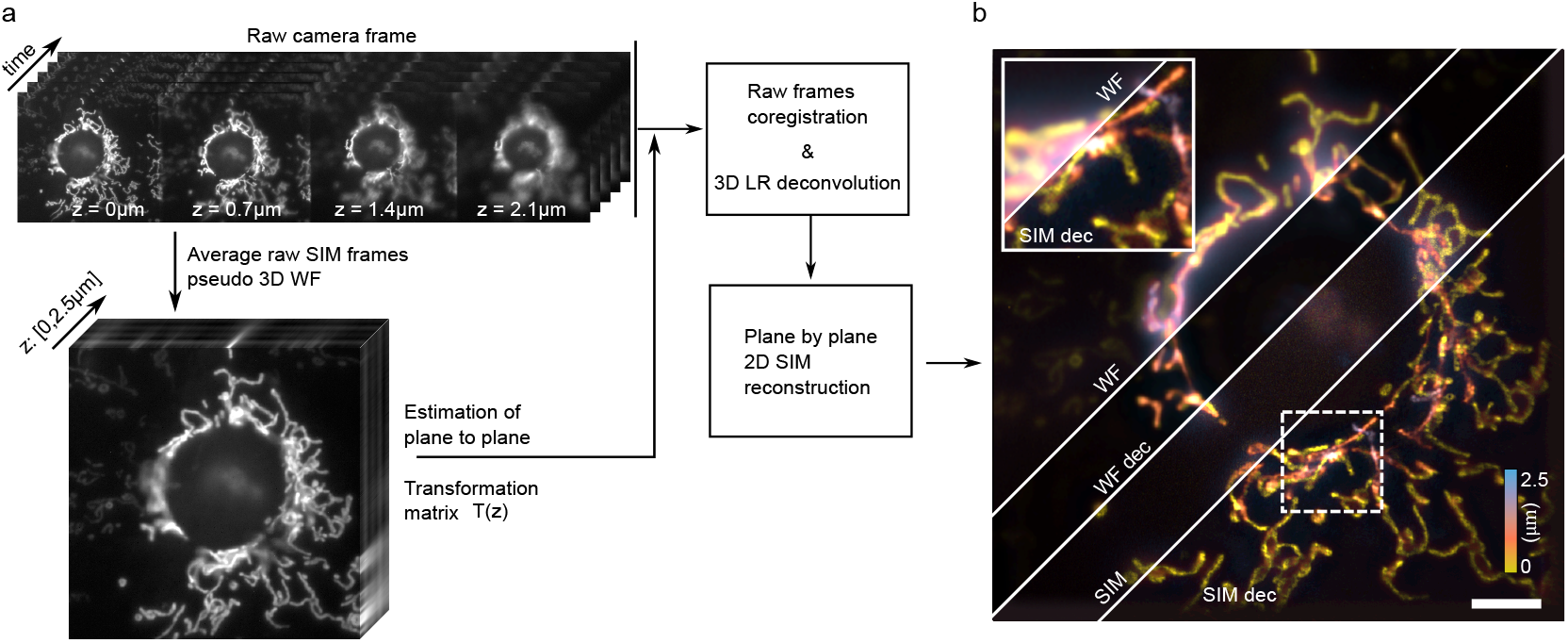
Data processing and SIM image reconstruction procedure. (a) The raw images are first averaged to produce a pseudo wide-field image. This image is then used to estimate the inter-plane transformation matrix *T* (*z*). This matrix is used to coregister the raw SIM images and reorder the data in x,y,z,t space. Then, the data is either averaged (WF), deconvolved in 3D (WF dec), reconstructed (SIM) or deconvolved and reconstructed (SIM dec). (b) Color coded maximum intensity z-projection of 3D images for all 4 modalities on a fixed COS-7 cells labelled with MitoTracker Orange. Inset shows a zoomed in comparison between WF and deconvolved SIM. Scale bar 5 *μm*.

Fig. 2a summarizes the flow of data. A key challenge is that the the 8 planes are all slightly shifted with respect to each other, due to unavoidable limitations in manufacturing. By cross-correlating two consecutive planes, we are able to recover the prism transformation matrix *T* (*z*) and perform a sub-pixel coregistration for all the raw SIM frames. We then average all the frames to form a pseudo-widefield image (WF), deconvolve the 3D WF using 10 iterations of Lucy-Richardson (LR) deconvolution (deconvlucy, Matlab 2017b), reconstruct the SIM image plane by plane (SIM) or deconvolve in 3D the raw SIM frames and reconstruct the SIM images plane by plane (SIM dec). Results are shown in Fig. 2b, where we encoded the depth information using a color-coded maximum z-projection. We used a custom 2D SIM reconstruction algorithm implemented in Matlab, following closely the work from [21]. For the 3D LR deconvolution, we used an experimentally acquired 3D PSF of the setup, obtained by imaging, localizing and averaging the 3D image of 15 sparsely distributed sub-diffraction fluorescent beads.

In Fig. 3, the time series of a second data set is shown. With an exposure time of 50 ms we achieved about 1.3 fps volumetric SIM imaging speed. We find (see Visualization 1) that this speed allows us to capture the 3D dynamics of the mitochondrial network. While the reconstructed sequence had to be bleach-corrected by normalizing the average of each frame, 74 time points (about 60 sec) could easily be acquired, without the need to resort to any advanced image reconstruction algorithms.

**Figure 3:**
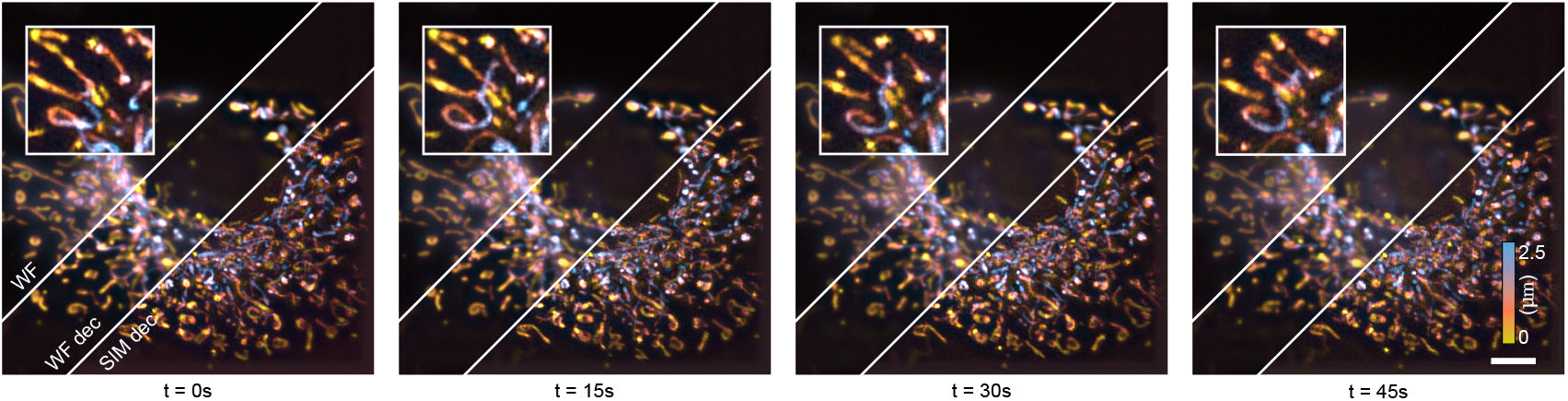
Time series of SR-SIM imaging data following mitochondrial dynamics (stained with MitoTracker Orange) in COS-7 cells, color coded maximum intensity z-projection of the deconvolved and SIM reconstructed 8 planes (see Fig. 2). The full series contains 74 time points, each acquired with 50 ms SR-SIM acquisition time to allow for motion-artifact-free imaging (see Visualization 1). The series has been renormalized in brightness to compensate for photo-bleaching, but no further processing beyond a standard SIM reconstruction had to be performed. Insets (FOV 7*x*7 *μm*^2^) show deconvolved SIM with pronounced mitochondria activity during the imaging. Scale bar 5 *μm*.

## 4 Conclusion

The results presented here demonstrate the first proof-of-concept of high-speed 2-beam multi-plane SR-SIM imaging of living cells using an image-splitting prism. They show that the combination of multi-plane image detection with fast 2-beam SR-SIM illumination indeed yields the desired high speed volumetric imaging. These results also indicate that this system is well suited to image mitochondrial dynamics, as both the necessary temporal and spatial resolution is reached.

Our work also provides several insights into current limitations and challenges. For example, the current SLM implementation encounters significant losses of the excitation light due to spurious diffraction, with a maximum of about 3 mW reaching the sample. Newer LCOS-based designs will allow for a 10× improvement in power management albeit at the cost of more complex timing requirements. In combination with more advanced algorithms taking full advantage of the 3D information for low signal to noise image reconstruction, we envision that 5-10 ms exposure times, and thus volumetric imaging within 50-100 ms will be possible. Very recent development into advanced denoising SR-SIM reconstruction algorithms[22] point to solutions that will allow for the use of significantly lower signal levels in the image reconstructions, and thus might allow us to push for even faster imaging speed. Advanced fluorescent dyes will allow for even more extended observation times. Novel, smart data-driven feedback loops should also be able to dynamically adapt the imaging speed depending on the observed dynamics. These approaches will all complement multi-plane video-rate SR-SIM imaging quite well.

In its current state, image-splitting multi-plane SR-SIM technology provides an early demonstration of what this technology will be able to achieve in more improved configurations. As the approaches and their implementations evolve, we believe they will provide an important tool for future high-speed super-resolution 3D imaging of living cells and organisms.

## Sample preparation and staining

#1.5 cover glass coverslips were cleaned with a piranha solution and coated with fibronectin (0.5 *μ*M/ml). Cells were grown in DMEM without phenol red medium, containing 10 % of fetal bovine serum. Mitochondria were stained with 100 nM MitoTracker Orange (Thermo Fisher) according to manufacturer-provided staining protocol for 30 min. Then cells were either fixed with 4% PFA or used for live cell imaging. For live-cell imaging, cells were washed twice with grown medium and imaged in PBS pH=7.4, which proved to reduce background fluorescence and short-term imaging was perform in a custom build incubator at 37 ° C.

## Acknowledgments

This project has received funding from the European Union’s Horizon 2020 research and innovation programme under the Marie Skłodowska-Curie grant agreements No. 752080 and No. 766181. T. Lasser and A. Radenovic acknowledges support from the Horizon 2020 framework program of the European Union via grant 686271. R.V.D.E. thanks the Research Foundation-Flanders for a doctoral fellowship, GBM-D5931-SB. P.D. acknowledges support by the European Research Council via ERC Starting Grant 714688 and from the Research Foundation-Flanders via grants G062616N, G0B8817N, G0A5817N, and VS.003.16N.

The authors declare that they have no conflicts of interest related to this article.

